# A Tool for Reliable Detection of Aflatoxin Biosynthetic Gene Clusters in Aflatoxigenic and Atoxigenic *Aspergillus flavus* Isolates

**DOI:** 10.1101/2021.03.31.437855

**Authors:** Alfred Mitema, Naser Aliye Feto, Sheila Okoth, Mohamed Suhail Rafudeen

## Abstract

Molecular techniques and phenotypic characterisation have been used to differentiate aflatoxigenic and atoxigenic *Aspergillus flavus* strains. However, there is a lack of a consistent and reliable tool for discrimination between these strains of *A. flavus*. Here we report, an optimised real-time qPCR-based tool for reliable differentiation between aflatoxigenic and atoxigenic strains of *A. flavus*. Accordingly, expression profiles and deletion patterns of genes responsible for aflatoxin production in five representative aflatoxigenic and atoxigenic *A. flavus* strains (KSM012, KSM014, HB021, HB026 and HB027) were examined using the optimised real-time qPCR tool. We observed that under induced conditions, *aflP, aflS, aflR* and *aflO* transcripts were the most upregulated genes across the tested isolates while *aflS* and *aflO* were always expressed in both induced and uninduced isolates. However, *aflR* and *aflP* did not give clear distinctions between non-toxin and toxin producing isolates. The deletion patterns were prominent for *aflD* and *aflR* whereas *alfO, aflS* and *aflP* had no deletions among the isolates. Significant variation in transcript abundance for *aflD, aflR* and *aflS* were observed for aflatoxigenic isolate KSM014 under induced and uninduced states. False detection of *aflD* gene transcript in atoxigenic strain KSM012 was evident in both induced and uninduced conditions. With the exception of KSM012, *aflP* gene did not exhibit significant variation in expression in the isolates between induced and uninduced conditions. One-way ANOVA and Post-test analysis for linear trends revealed that aflatoxin biosynthetic cluster genes show significant (P < 0.05) differences between atoxigenic and aflatoxigenic isolates. Our optimized qPCR-based tool reliably discriminated between aflatoxigenic and atoxigenic *A. flavus* isolates and could complement existing detection methods.

## Introduction

Certain fungi of the *Aspergillus* genus produce secondary metabolites termed aflatoxins, which are a class of naturally-occurring mycotoxins (1). About 200 species of *Aspergillus* have been identified, of which 16 have been found to produce aflatoxins that are detrimental to both human and animals (1,2). A number of *Aspergillus* species such as *Aspergillus flavus, Aspergillus bombycis, Aspergillus nomius* and *Aspergillus niger* produce aflatoxins with high carcinogenic activity (3,4). Aflatoxin contamination has been detected in maize, beans, cottonseed, peanuts and other grain crops (5,6). The contamination not only results in reduced crop value but can cause health problems in both humans and animals that consume contaminated crops and feeds (7).

Although aflatoxin-producing *Aspergillus* species are found worldwide, they are of greater concern in underdeveloped countries which lack appropriate infrastructure, management tools and resources required to prevent, control or monitor their impact on the wider community (1). Environmental conditions: unseasonal rains during harvesting, increased temperatures and moisture promotes fungal pathogen proliferation and mycotoxin production (8,9). Additionally, increased risks of mycotoxin production and fungal growth is facilitated by improper harvesting, poor storage facilities and sub-optimal temperatures during processing and marketing. These environmental conditions and problems associated with food production and storage are common in most parts of sub-Saharan Africa, where to date, the largest poisoning of mycotoxin epidemic has been reported (10,11).

The classification and identification of *Aspergillus* spp. based on phenotype is complemented by molecular and chemotaxonomic characterisation (12,13). Phenotypic characterisation of *A. flavus* isolates may identify an isolate as potentially aflatoxigenic, but this is neither definitive nor precise (14,15). Molecular techniques, such as Reverse Transcriptase Polymerase Chain Reaction (RT-PCR) have also been used to differentiate aflatoxigenic from atoxigenic *A. flavus* strains, using the expression of regulatory and structural aflatoxin pathway genes as markers for aflatoxin production (16,17). Despite the complexity of the aflatoxin pathway involving at least 25 structural and two regulatory genes (18), some studies have found good agreement between gene expression and aflatoxin production (17). Scherm et al. (17), reported the expression profiles of the genes *aflO, aflP* and *aflD* were linked with the *A. flavus* strains capability for aflatoxins production.

In *A. flavus*, expression of *nor-1* (*aflD*), the gene that encodes for an enzyme meant to catalyse the conversion of the first stable biosynthesis of aflatoxins, (Fig. S1) norsolorinic acid, to averantin (19,20) is the main structural gene in biosynthetic pathway of aflatoxins. Previous studies established the transcription *of aflD* as a better marker for discrimination between atoxigenic and aflatoxigenic strains (21,22). In contrast, *aflR* transcription, which expresses a regulator which is capable of activating majority of structural genes in the aflatoxin biosynthetic pathway had poor discriminatory power (23).

The *aflS* gene codes for regulatory protein, laeA (24) whose function is not confirmed though, (25) postulated that the interaction between *aflR* with laeA may support binding of DNA by the former. Du et al. and Schmidt-heydt et al. (26,27), also observed that *aflS* may have the ability in modulation of *aflR* activity and that environmental factors may influence the ratio of *aflR* to *aflS*. According to (28), in the case of some *Aspergillus* spp., this modulation may have shifted the affinity of *aflR* towards the G-group-specific genes, *ordA, cypA* and *nadA*, and activated their expression under environmental factors.

The potential molecular mechanisms that lead to the loss of aflatoxin production are diverse, and for most atoxigenic *A. flavus* strains, the specific genetic mechanisms resulting in atoxigenicity have not been elucidated in great detail (29).

In the current study, the expression profiles of genes responsible for aflatoxin production in five aflatoxigenic and atoxigenic *A. flavus* strains (KSM012, KSM014, HB021, HB026 and HB027) were examined by real-time qPCR. The isolates were chosen from four different climatic regions of Kenya based on certain morphological and phenotypic characteristics (30).

## Results

### RNA extraction

The RNA extracted from *A. flavus* strains cultivated on YES and YEP medium had acceptable 260/280 ratios (Table S1). The RNA was of good quality and integrity as seen by clear ribosomal RNA bands (Fig.1a). Additionally, there was successful cDNA synthesis, indicated by a continuous smooth cDNA smear (Fig.1b).

**Figure 1.**
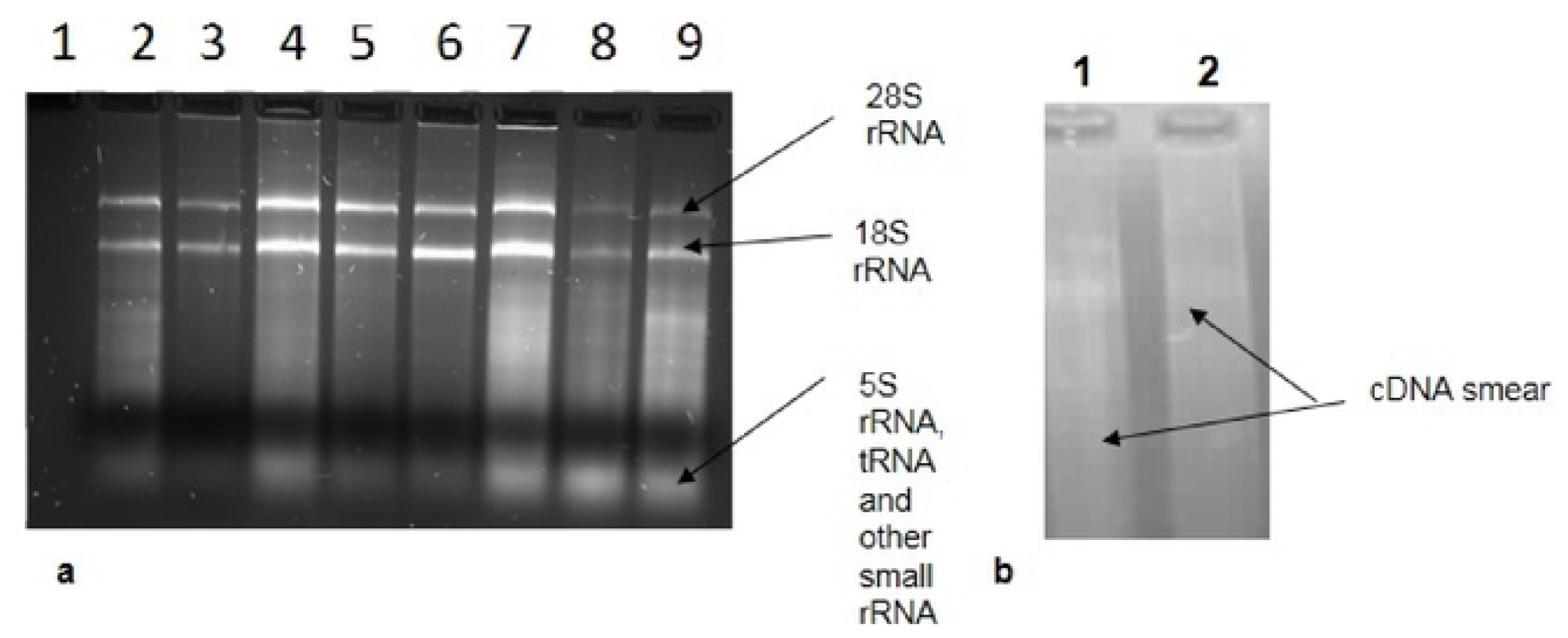
Gel electrophoresis of RNA extracted from selected *Aspergillus flavus* strains assessed on a 1.2 % agarose/EtBr gel at 80V for 45 min showing; (**a**). Total RNA; Lane: **1**. No template (control); Lanes **2**. NC05; **3**. KSM012; **4**. KSM014; **5**. HB021; **6**. HB025; **7**. HB26; **8**. HB027; **9**. HB028; (**b)**. cDNA synthesis (Lane: **1**. KSM012 and **2**. KSM014). [(NC: Nandi county, Kenya); (KSM: Kisumu, Kenya); (HB: Homa Bay, Kenya); EtBr: ethidium bromide; cDNA: complementary deoxyribonucleic acid; RNA: ribonucleic acid].

### Aflatoxin biosynthetic gene cluster and amplicon sizes

We measured the expression profiles of regulatory and structural genes responsible for aflatoxin production in selected aflatoxigenic and atoxigenic *A. flavus* strains using real-time qPCR. Five specific isolates of *A. flavus* previously deposited (31) in NCBI GenBank: (KSM012_MG385137, KSM014_MG385138, HB021_MG430317, HB026_MG430319 and HB027_MG430322) were chosen based on their sample location in Kenya together with their S and L-morphotype and ability to produce sclerotia. We used six sets of primers (Table 1) to detect the presence or absence of aflatoxin genes in the induced or un-induced isolates. The cDNA from the respective *A. flavus* isolates from both induced and uninduced states were subject to qPCR with primers specific for genes in the aflatoxin gene cluster (Fig.2; Table 1). Amplicon sizes exhibited by *A. flavus* isolates were 120, 116, 117, 123, 168, and 118 bp for *aflO, aflD, aflS, aflP, aflR* and *β-Tub*, respectively (Fig.2a-f).

**Table 1.** Different gene expression profiles or deletion patterns exhibited by *Aspergillus flavus* strains.*

| Isolates | County | Strain | UV 365nm | Sclerotia |  | Status | Mean normalised Ct value ratios (log<sub>10</sub>) |  |  |  |  |
| --- | --- | --- | --- | --- | --- | --- | --- | --- | --- | --- | --- |
|  |  |  |  | S | L |  | *aflP* | *aflD* | *aflO* | *aflR* | *aflS* |
| 12 | KSM | atoxigenic | green | - | - | induced | 0.150±0.017 | 0.117±0.008 | 0.059±0.012 | -0.007±0.010 | 0.034 ± 0.010 |
|  |  |  |  |  |  | Uninduced | 0.093±0.032 | 0.099±0.003 | -0.067±0.004 | -0.027±0.006 | 0.033 ± 0.004 |
| 14 | KSM | aflatoxigenic | blue | ++ | - | induced | 0.254±0.029 | 0.278±0.007 | 0.139±0.011 | 0.144±0.024 | 0.233 ± 0.008 |
|  |  |  |  |  |  | Uninduced | 0.218±0.065 | DL | 0.181±0.010 | DL | 0.259 ± 0.007 |
| 21 | HB | afl/atoxigenic | blue/green | - | - | induced | 0.177±0.006 | DL | 0.113±0.003 | DL | 0.153 ± 0.011 |
|  |  |  |  |  |  | Uninduced | 0.112±0.026 | DL | -0.066±0.003 | DL | 0.024 ± 0.004 |
| 26 | HB | afl/atoxigenic | blue/green | - | ++ | induced | 0.089±0.007 | DL | 0.029±0.012 | DL | 0.069 ± 0.004 |
|  |  |  |  |  |  | Uninduced | 0.137±0.013 | DL | -0.051±0.004 | DL | 0.012 ± 0.006 |
| 27 | HB | afl/atoxigenic | blue/green | - | ++ | induced | 0.130±0.022 | DL | 0.078±0.056 | DL | 0.175 ± 0.004 |
|  |  |  |  |  |  | Uninduced | 0.109±0.008 | DL | -0.051±0.004 | DL | 0.036 ± 0.010 |

\* Threshold cycle value ratios, mean ± standard deviation error based on three biological replicates.
**DL:** deletion
- or ++: absence or presence of sclerotia
**S** or **L**-morphotype of sclerotia
**KSM:** Kisumu
**HB:** Homa Bay

**Figure 2.**
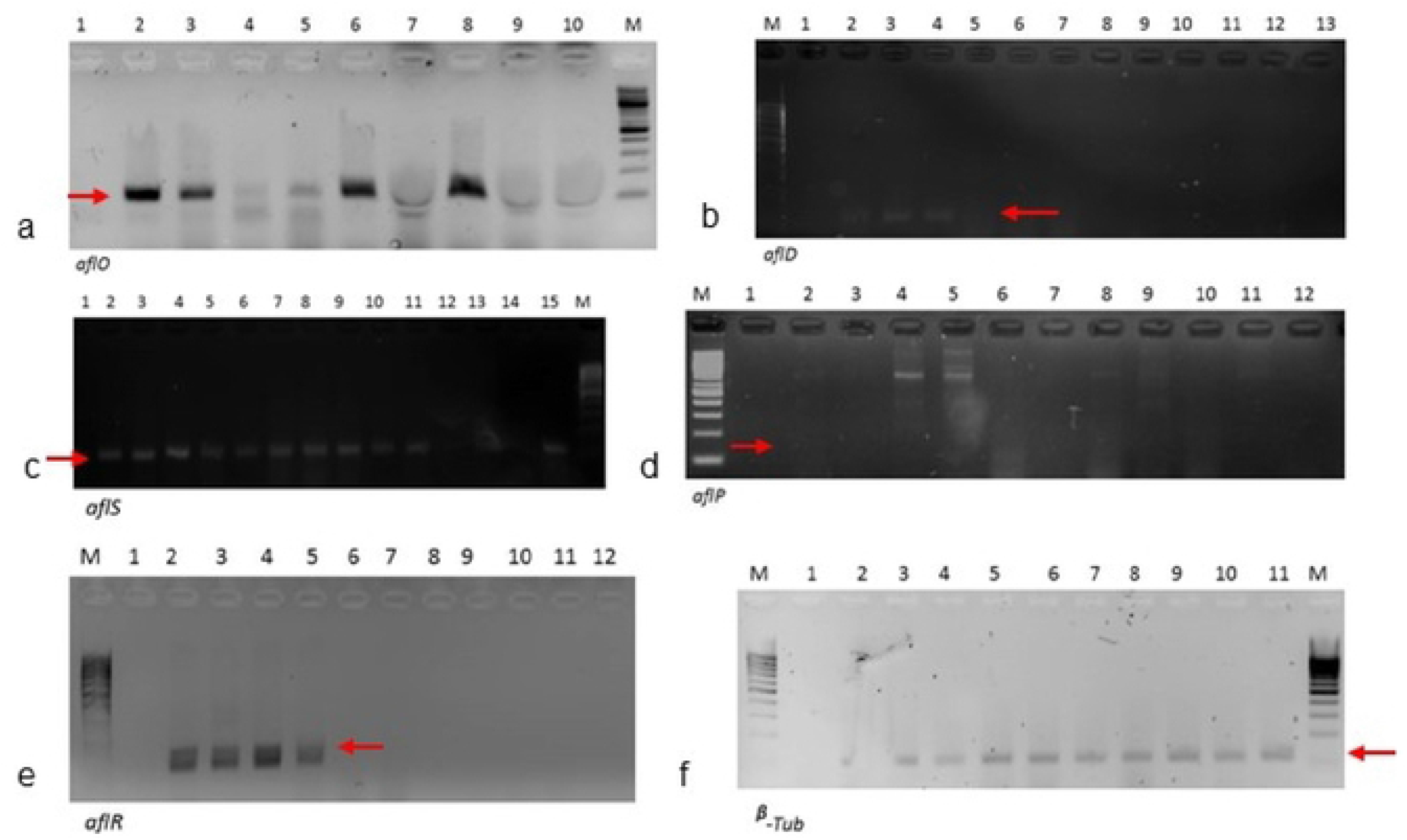
Gel electrophoresis of induced and uninduced isolates of *A. flavus* assessed showing qPCR product amplicon sizes of different genes run at 80V for 45 min on 2% agarose/EtBr gel displaying various deletion patterns. **M**: 100 bp low range molecular marker; ***aflO***: Lane **1**: NTC; **2**. Unind27; **3**. Ind27; **4**. Unind26; **5**. Ind26; **6**. Unind21; **7**. Ind21; **8**. Unind14; **9**. Ind14; **10**. Unind12; ***aflD* and *aflR*;** Lane **1**. NTC; **2**. Ind12; **3**. Unind12; **4**. Ind14; **5**. Unind14; **6**. Ind21; **7**. Unind21; **8**. Ind26; **9**. Unind26; **10**. Ind27; **11**. Unind27; **12**. noRT; **13**. noRT; ***aflS***: Lane **1**: NTC; **2**. Unind27; **3**. Ind27; **4**. Unind26; **5**. Ind26; **6**. Unind21; **7**. Ind21; **8**. Unind14; **9**. Ind14; **10**. Unind12; **11**. Ind12; **12**. NTC; **13**. Pldstd3; **14**. Pldstd2; **15**. Pldstd1; ***aflP***: **1**. Pldstd1; **2**. Unind27; **3**. Ind27; **4**. Unind26; **5**. Ind26; **6**. Unind21; **7**. Ind21; **8**. Unind14; **9**. Ind14; **10**. Unind12; **11**. Ind12; **12**. NTC; **β-Tub**: Lane **1**: NTC; **2**. Unind27; **3**. Ind27; **4**. Unind26; **5**. Ind26; **6**. Unind21; **7**. Ind21; **8**. Unind14; **9**. Ind14; **10**. Unind12; **11**. Ind12. The isolates were cultured on inducing media (YES) and non-inducing media (YEP) (Unind: Uninduced; Ind: induced and Pldstd: Pooled standard).

### qPCR and primer efficiency analysis

The amplicons generated by qPCR showed that the primers (Table 1) were specific, appropriately designed, and suitable for studying *A. flavus* and aflatoxin genes (Tables 1; 2). The *aflR* and *aflD* had unique expression profiles and deletion patterns (Table 1; Fig.2). The standard melt curves exhibited statistical linear regression values and efficiency range (Table 2).

**Table 2.**
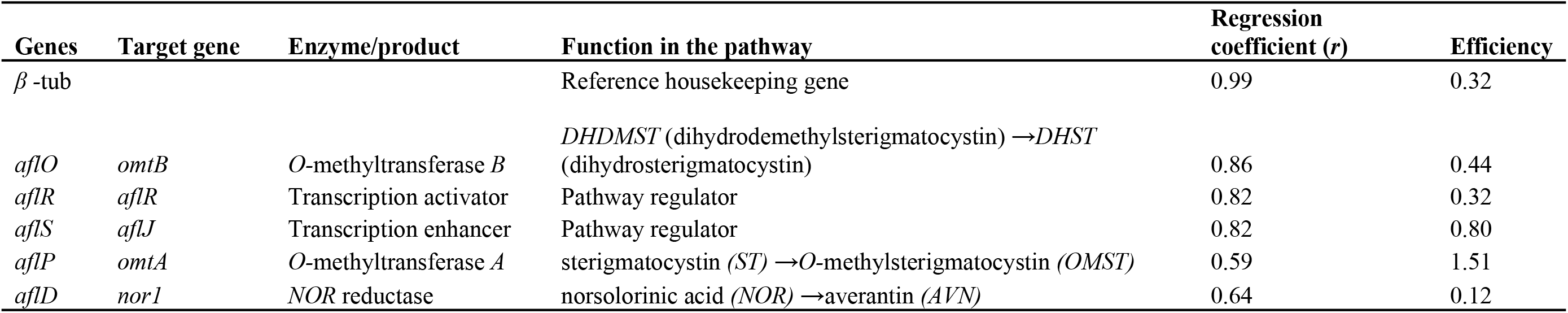
Clustered aflatoxin biosynthesis pathway genes showing enzymes involved, functions, statistical linear regression and efficiency.

When uninduced, *aflD* and *aflR* exhibits the deletion pattern for *A. flavus* strains HB021, HB026, HB027, and KSM014 (Table 2, Figs. 2; 3a; 4c). These genes were expressed by KSM012 under induced and uninduced conditions and KSM014 under induced conditions (Table 2). In contrast, the *aflP, aflS* and *aflO* genes were expressed by all five *A. flavus* strains under both induced and uninduced conditions. From the observation, it can be suggested that these *aflP, aflS* and *aflO* expression are not ideal indicators for determining whether a strain can synthesize aflatoxins.

**Figure 3.**
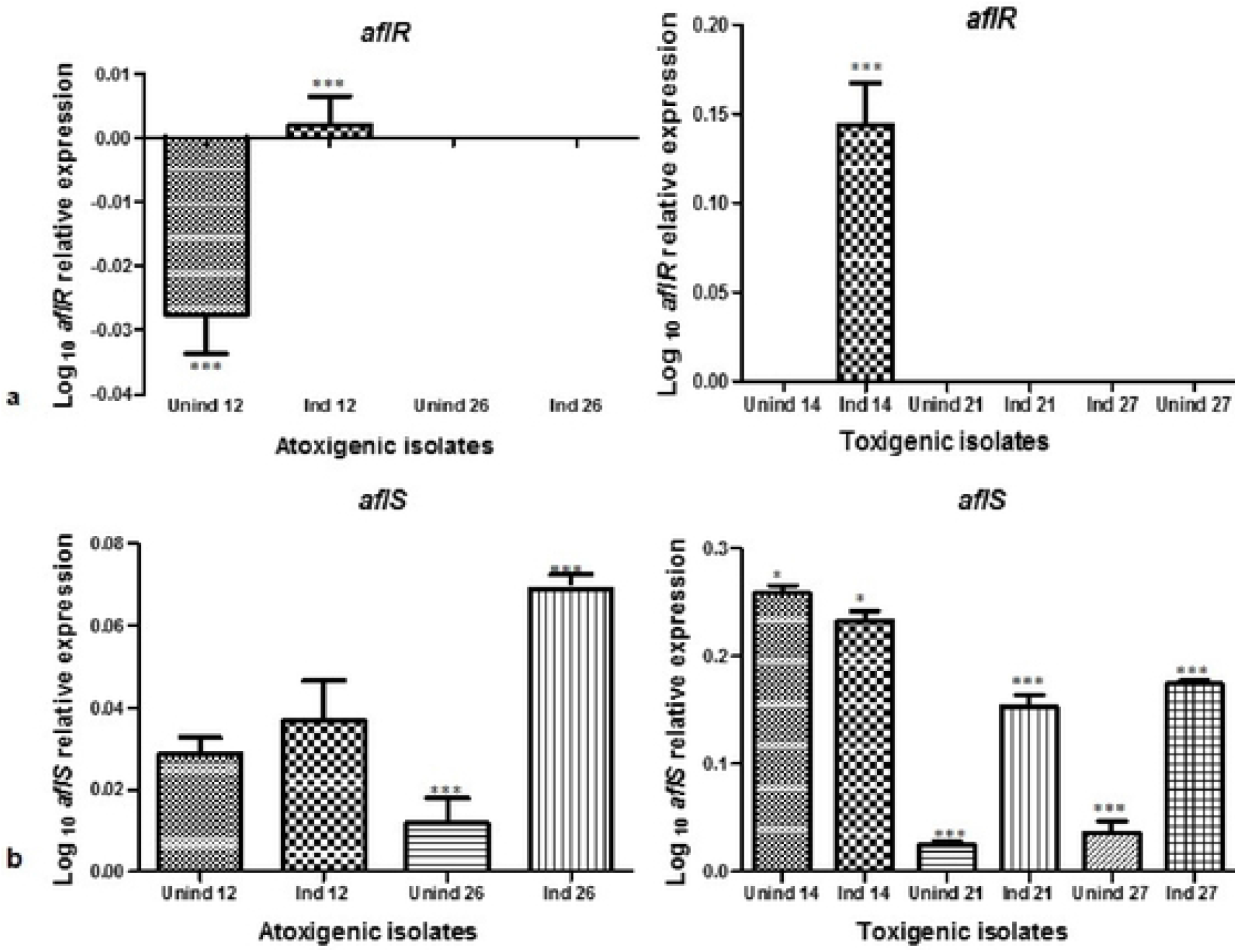
Expression profiles for *aflR* and *aflS*. **a**. Significant difference in expression were noted in *aflR* for isolates 12 and 14 (both induced), with isolate 14 upregulated significantly. Isolates HB021, HBO26 and HB027 displayed no significant expression. Uninduced isolate KSM012 was significantly down regulated. **b**. *aflS* expression profiles. The expression values were normalised, and log transformed (log_10_). Asterisks and the error bars shows significance variance and standard mean deviations (*n* = 3), 1-way ANOVA and Tukey’s Multiple Comparison Test (P < 0.05). (KSM: Kisumu; HB: Homa Bay; Unind: uninduced; Ind: induced; 12-27: isolates).

**Figure 4.**
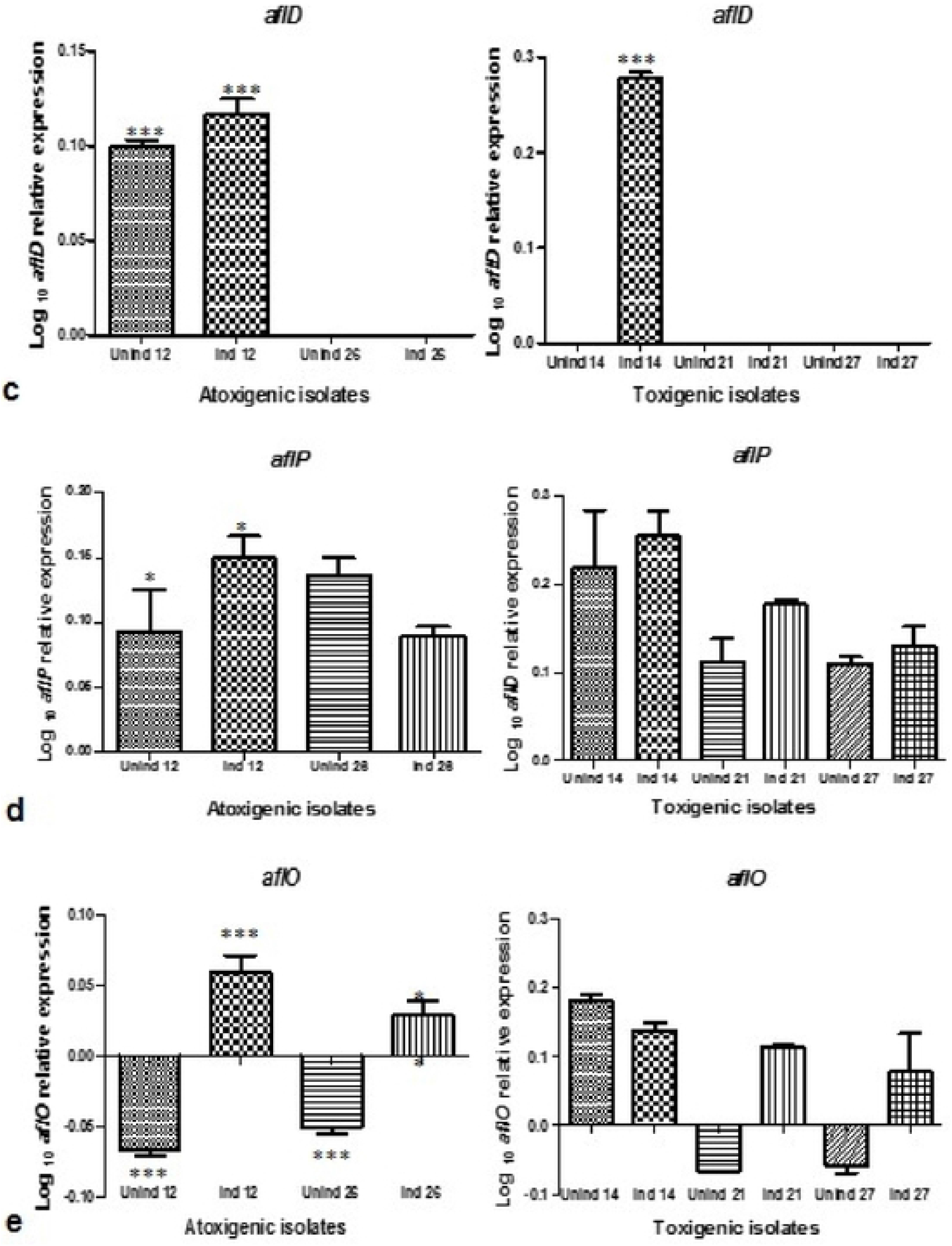
Aflatoxin biosynthetic gene cluster expression profiles. **c**. *aflD* expression profiles for *Aspergillus flavus* KSM012 and KSM014, with isolate KSM014 highly expressed. No expression was observed for isolates HB021, HB026 and HB027. **d**. *aflP* expression, which was statistically significant only for isolate KSM012. **e**. *aflO* expression profiles. There were significant difference for the atoxigenic isolates but not for the toxigenic isolates. Expression values were normalised, and log transformed (log_10_). [Asterisks and the error bars show significance variance and standard deviations of the mean, 1-way ANOVA and Tukey’s Multiple Comparison Test (P < 0.05)]. (KSM: Kisumu; HB: Homa Bay; Unind: uninduced; Ind: induced; 12-27: isolates).Significant decreases in *aflR* transcripts (P < 0.05) were found for strains KSM012, HB021, HB026 and HB027 (Fig. 2.3a). In *A. flavus* KSM014, an aflatoxin producing strain, *aflR* transcript abundance increased under induced and decreased under uninduced conditions (Fig.3a). Thus, *aflR* could be a marker to differentiate toxin and non-toxin producers. When comparing the induced isolates, isolate HB026, had higher levels of expression of *aflS* and *aflO*. KSM012 and KSM014 had higher *aflD* and *aflR* transcript levels in induced than in uninduced cultures (Figs.3; 4; Table 2). *AflR* expression decreased significantly in uninduced isolate KSM012 (Fig.3a).

### *A*flatoxin biosynthesis pathway gene cluster expression profiles

Expression profiles of the structural genes (*aflO, aflP, aflD*) and regulator genes (*aflS, aflR*) were analysed. One-way analysis of variance (1-way ANOVA) and a Post-test for linear trend confirmed significant differences (P < 0.05) between atoxigenic and aflatoxigenic isolates for some of the isolates (Fig.3; 4).

Tukey’s Multiple Comparison Test (TMCT) revealed significant variances for different biosynthetic genes (Figs.3; 4).

## Discussion

The designed primers were suitable for the current study and our findings (Tables 1; 2; 3 and Fig.2) were similar to previously reported studies (27,32) who found that decreased *aflD* (*nor-1*) expression was correlated with a subsequent decrease in aflatoxin production in an aflatoxigenic strain. The *aflD* is an important gene in the biosynthesis pathway in both *A. flavus* and *A. parasiticus* (19,22). The inhibition or disruption of *alfD* gene expression results in the accumulation of intermediates in aflatoxin biosynthesis such as norsolorinic acid and prevents the synthesis of aflatoxins and any other intermediates in the pathway (33). Similarly (27,32) observed that the disruption of aflatoxin production occurs if specific inhibitor (enzyme) blocks the biosynthesis pathway. Scherm et al. (17), also found that a reduction in aflatoxin level was accompanied by a decrease in *aflD* transcripts. Furthermore they found that the expression profiles of *aflD* (*nor-1*), *aflO* (*omtB*) and *aflP* (*omtA*) were consistently correlated with the production of aflatoxins, whereas the expression of *aflS* (*aflJ*) and *aflR* were not.

Data analysis from Table 2 showed that *aflP, aflS* and *aflO* gene expression is not essential for determining whether a strain can synthesize aflatoxins. Cary & Ehrlich (23), established that *aflR* is a regulatory gene of the pathway intricated in transcription and activation of majority of the aflatoxin biosynthetic structural genes. Schmidt-Heydt et al. (24), found that the ratio between regulatory genes *aflS* and *aflR* that is activated by environmental factors. Schmidt-heydt et al. and Abdel-Hadi et al. (22,27) demonstrated the potential transcription of *aflD* as a marker to distinguish aflatoxin producing and non-producing isolates from peanut, while *aflR* expression could not be used to differentiate these phenotypes.

In the current study, an aflatoxin producing strain KSM014, consistently had higher transcript levels than the other isolates across all the studied genes (Figs.3; 4). For *aflR* and *aflD*, there was no significant increase in transcript abundance in isolate KSM014 (uninduced state). The *aflO* and *aflS* genes were expressed at higher levels in the uninduced state than in the induced (Figs. 3b; 4e). This observation suggests that there could be a relationship between *aflO* (structural gene) and *aflS* (regulatory gene). Furthermore, *aflO* expression was significantly lower under induced conditions than uninduced conditions (Fig.4e). *aflO* expression decreased in uninduced isolates of KSM012, HB021, HB026 and HB027 but increased for isolate KSM014 (Fig.4e).

Expression of the *aflP* gene did not vary significantly between induced and uninduced isolates KSM014, HB021, HB026 and isolate HB027 according to 1-Way ANOVA and TMCT test (Fig.4d). *aflP* expression was significantly higher for isolate KSM012 in both induced and un-induced states. Expression in the uninduced isolate, HB026 was higher than in the induced isolate, but the difference was not significant based on a TMCT test. This observation suggests that *aflP* is not a promising marker that can be used in discrimination of aflatoxigenic and atoxigenic isolates (Fig.4d). *aflD, aflR* and *aflS* all differed significantly in expression in KSM014 between induced and uninduced states (Figs.3; 4).

The *aflD* gene transcript was detected in both induced and uninduced isolate KSM012, which is atoxigenic (Fig.4c; Table 2). False positive and negative transcription signals have been previously observed (34). Biosynthesis of aflatoxin in *Aspergillus* is regulated by sophisticated pattern of negative and positive transcriptional regulatory factors, that are sensitive to both external and internal stimuli (34). Chiou et al. (2002), postulated that chromosomal position of important genes may take part in gene expression of aflatoxin, and that, PCR based screening techniques involving genes in the aflatoxin biosynthesis pathway may flop in distinguishing atoxigenic and aflatoxigenic strains from one another.

Chang et al. (36), found that all *A. flavus* strains, both atoxigenic and aflatoxigenic carry *aflD, aflP* and *aflR*. This result indicates that the absence of production of aflatoxin in some isolates is due to degeneration of the entire cluster genome. The loss of production of AFB1 and AFB2 by many atoxigenic *A. flavus* isolates is not due to large deletions, but could also arise from point mutations (36).

Moreover, real time-PCR outcomes have indicated that transcription of *aflD* may be used as an indicator in discrimination of aflatoxin and non-aflatoxin producers, while *aflR* and *aflP* did not discriminate aflatoxigenic and non-aflatoxigenic strains (17). Similarly, they further observed that *nor-1*, gave good correlation of aflatoxin production and gene expression on non-inducing and inducing media. Their findings contrasts to that of (37), wherein the study involved expression analyses of *aflO* and *aflD* genes, in *Aspergillus* section *Flavi* isolates (31), native almonds from Portuguese. Their conclusion was that expression *of aflD* was not a promising marker that could be used in differentiation of non-aflatoxigenic and aflatoxigenic isolates. There was only one almond isolate in their study that gave false positive transcript. We have also observed in the current study that *aflD* and *aflR* transcripts were not consistent in giving a clear distinction between aflatoxigenic and atoxigenic strains, which is in line with a related finding by (37). This could be due to either point mutations or large deletions in the aflatoxin genome cluster involved in aflatoxin production (Fig.3a; 4c).

Based on our results, *aflO* might be considered as a marker for distinguishing atoxigenic and aflatoxigenic strains (Fig.4e; Table 2). We used a more sensitive real-time RT-qPCR approach compared to other studies. This involved different marker genes (MEP for maize, *β-tub* and *Ef1a* for *A. flavus*) and using SYBR green fluorescence to detect expression. Scherm et al. and Rodrigues et al. (17,37) who obtained contradictory results to each other used the less sensitive RT-PCR method and only the *A. flavus β-tub* gene as a marker. Due to the inconsistency of results across research groups, further studies are needed to identify a molecular marker suitable for differentiating atoxigenic and aflatoxigenic strains. The qPCR protocol used in this study gave promising results and should be recommended in future research especially with *A. flavus* strains. Criseo et al. (38) also used molecular techniques to distinguish non-aflatoxigenic and aflatoxigenic strains of *A. flavus* through the correlation with the presence or absence of one or several genes in the biosynthesis pathway of aflatoxin.

Studies differentiating aflatoxigenic and atoxigenic strains have often relied on monitoring aflatoxin gene expression with reverse transcription and real-time PCR methodologies (16,39). It is, thus, understandable that lack of a tool/protocol for reliable discrimination between toxigenic and atoxigenic strains of *A. flavus* isolates have not yet been successfully established. Furthermore, one should be aware that some genes are not exclusive to the aflatoxin biosynthetic pathway, which could create false-positives from sterigmatocystin producing fungi (40).

In conclusion, many enzymatic steps are involved in the aflatoxin biosynthesis pathway. Expression level measurements of the genes that codes for the enzymes or the absence/presence of these genes could provide information on whether a strain is aflatoxigenic or not. However, there is disparity in identifying a suitable gene indicator for production of aflatoxin and measuring a strains ability to produce aflatoxin remains the only reliable method. High genetic similarity between species of *Aspergillus* and high levels of intraspecific genetic variation have hindered reliable identification of molecular markers capable of consistently differentiating various strains and their aflatoxigenic potential and makes the task challenging.

We found that certain genes in the aflatoxin biosynthetic pathway are expressed more highly than others in the same pathway. Accordingly, expression increased the most for *aflP, aflS, aflR* and *aflO*. The modifications and optimisation of the RT-qPCR method used in this study enabled better discrimination of the isolates with respect to their possible toxigenicity or non-toxigenicity. We recommend this approach with suggested modifications for future work on reliable discriminating between toxigenic and atoxigenic strains of *A. flavus* isolates.

## Materials and methods

### *Aspergillus flavus* cultures

Fungal cultures from previous studies (30) were consistently maintained on new PDA plates (Sigma Aldrich, USA) complemented with chloramphenicol for inhibition of bacterial growth. Mono-conidial bacterial cultures were cultivated by taking a small quantity of mycelium and transferring into 1000 µl of sterile water. One hundred microliters of the inoculum were then transferred onto the water agar (WA) plate, swirled for a few seconds with conidial suspension and liquid culture discarded off the plate, thereafter, the plate was incubated for 12 hrs at RT. Mono-conidial cultures were assessed under stereomicroscope (Nikon Corp., Japan), excised, relocated onto new PDA plates and incubated at 25 °C in the dark. For aflatoxin induction, each *Aspergillus* isolate was grown in non-inducing medium yeast extract peptone (YEP) (Sigma Aldrich, USA) and aflatoxin-inducing medium yeast extract agar (YES) (Sigma Aldrich, USA) plates and incubated for 5 days at 25 °C prior to ribonucleic acid (RNA) extraction.

### Total RNA extraction

RNA was extracted from each *A. flavus* isolates grown on medium (YEP and YES) respectively. Mycelia were scrapped off the plates, flash frozen and ground in liquid nitrogen in a sterile pestle and mortar. Approximately 200-300 mg of ground mycelium was flooded with TrizoL^®^, 750 µl (Sigma Aldrich, USA) in 2 ml tubes containing 0.3 mm diameter glass beads (Merck KGaA, Darmstadt, Germany). The mixture was vortexed, thereafter incubated for 10 min at room temperature (RT) range 21-23 °C. Chloroform (200 µl) (Sigma Aldrich, USA) was added to the homogenized sample and mixed gently for 1 min. The tubes were incubated for 5 min at RT, followed by centrifugation for 15 min (14000 x g) at 4 °C (Beckman Coulter, Inc USA). The aqueous phase was transferred to a fresh 1.5 ml tube and the organic phase was kept for deoxyribonucleic acid (DNA) and protein extraction at -80 °C. Isopropanol (500 µl) (Sigma Aldrich, USA) was added to the aqueous phase and incubated for 10 min at RT to allow RNA precipitation. The mixture was then centrifuged for 10 min (14000 x g) at 4 °C and the supernatant discarded. The pellets were washed in 1 ml of cold 75 % ethanol (Sigma Aldrich, USA) for 1 min, then centrifuged for 5 min (14000 x g) at 4 °C. The supernatant was discarded, and the RNA pellet air-dried for 10 min at RT. The pellets were re-suspended in100 µl DEPC water, incubated at 55 °C for 10-15 min and then stored at -80 °C until analysed.

The RNA extracts were then treated with deoxyribonuclease I (DNase I; New England Biolabs, USA) to digest and remove any genomic DNA contaminant as per the manufacturer’s instruction. The sample was then purified using the Zymo Research Fungal/Bacterial RNA Miniprep Kit (Inqaba Biotec, South Africa) according to the manufacturer’s instructions. The RNA was measured with a Nano Drop ND-1000 spectrophotometer (Nano Drop Technologies, USA) and the integrity was assessed on a 1.2 % agarose/EtBr gel.

### First strand cDNA synthesis

Circa 500 ng of RNA was used for cDNA synthesis. The reaction was performed in triplicate using M-MLV Reverse Transcriptase Kit (Promega, USA) according to the manufacturer’s instructions with modifications. Briefly, a ratio of 1:1 instead of 1:10 random hexamers to Oligo (dT) was selected for cDNA synthesis from the fungal RNA in order to increase sensitivity. The primer mix (reaction volumes of 10 µl consisted of 5 µl nuclease free water, 1 µl 50 ng/ml Random hexamer, 0.1 µl of 500 ng/ml Oligo dT and 500 ng RNA) was incubated at 70 °C for 5 min and immediately cooled on ice for 5 min, then briefly centrifuged. The annealed primer mix was added to the master mix (reaction volumes of 14.5 µl nuclease free water, 10 µl 5 X M-MLV reaction buffer, 2 µl M-MLV-RT point mutant, 2.5 µl dNTP Mix and 1 µl inhibitor) in the ratio of 1:3, gently mixed, and then aliquoted onto PCR tubes. The reaction conditions consisted of four cycles of 25 °C for 20 min, 37 °C for 40 min, 42 °C for 90 min, followed by an incubation at 70 °C for 15 min and a final step at 4 °C for 1 min. To assess for successful cDNA synthesis, samples were initially run on a 1.2 % (w/v) agarose/EtBr gel at 120V for 5 min, thereafter, at 80 V for 45 min and visualised as previously described (31). The synthesised samples were combined and stored at -20 °C for subsequent use and at -80 °C for later analyses.

### qPCR and primer design

To detect the presence or absence of aflatoxin genes in the induced or un-induced isolates, six sets of primers (Table 3) for one reference gene (*β-*tubulin) and five genes of interests (structural and regulatory) were designed and assessed as previously described (30,41). The PCR and melt curve analysis were used to identify both specific and non-specific amplification.

**Table 3.**
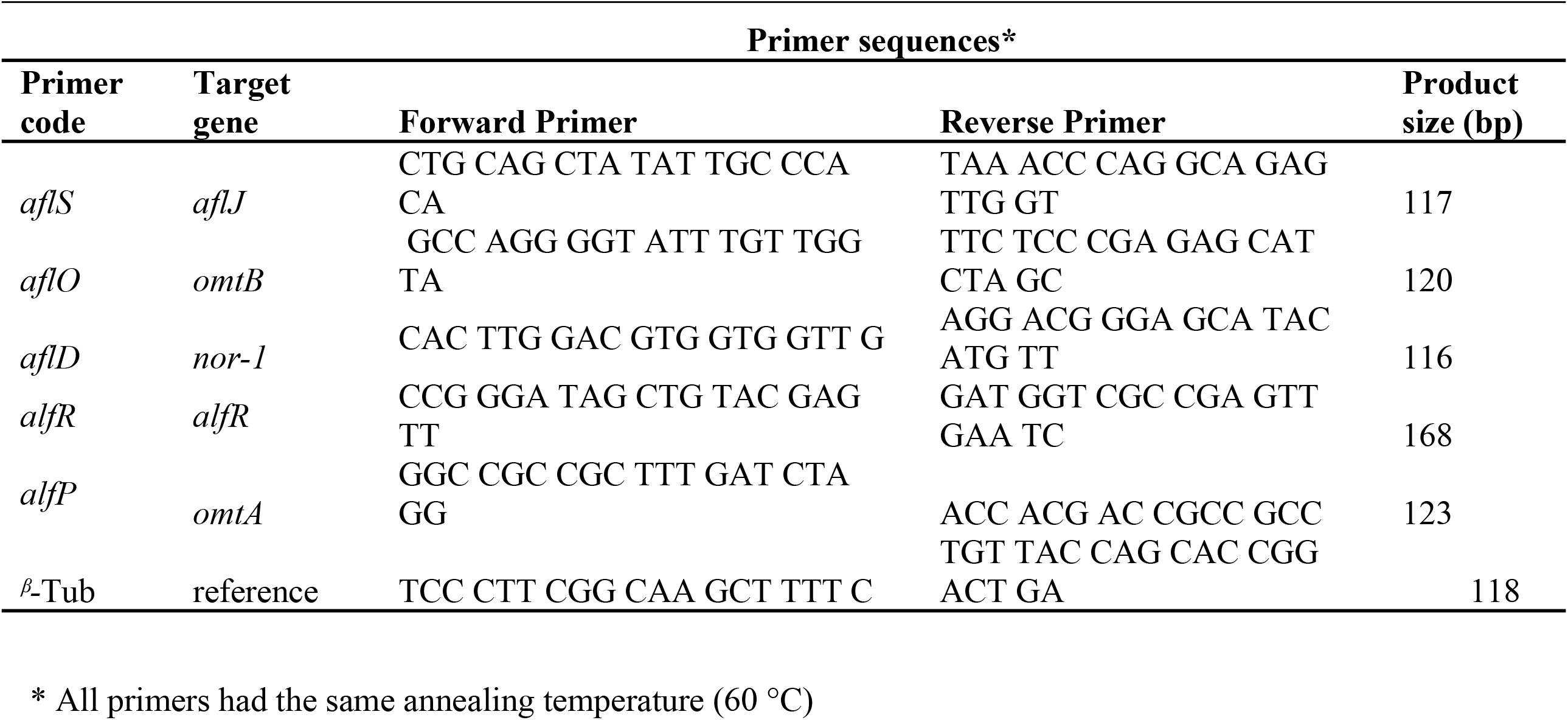
List of primers and the corresponding target genes used in the study.

### qPCR efficiency

The expression profiles and analysis of the genes were investigated using Rotor Gene 6000 2 plex HRM (Corbett Life Science Research, Australia). Serial dilutions of pooled cDNA (10-fold) from induced and uninduced isolates were used to generate standard curves. The Kapa SYBR Fast Kit (Kapa BioSystems, South Africa) master mix containing reaction buffer, heat-activated DNA polymerase, dNTPs and a working concentration of 3 mM MgCl_2_ were used for each qPCR reaction. The reaction consisted of a final concentration of 1 X Kapa SYBR green, 10 μM gene specific primers (0.4 µl), 1 µl of cDNA and nuclease free water to a total volume of 20 µl. The primer sets (Table 1) were used in separate reactions. Each dilution-point reaction was performed in triplicate along with a no-reverse transcription control, and a no-template control (NTC) in each real-time run. Amplification was carried out under the following conditions: 95 °C for 10 min; and 35 cycles of 95 °C for 3 s, 60 °C for 20 s, and 72 °C for 1 s.

### Expression stability analysis of aflatoxin biosynthetic genes

The qPCR reaction mixes and conditions were set up as described previously (30). To minimise variations between qPCR runs, all the reactions containing one primer pair were performed in a single run. The average expression levels were calculated from three technical repeats and by importing the relative standard curve into each run. Relative gene expression was determined by the amplification threshold in the exponential phase of the PCR, identifying the threshold cycle (Ct) value and comparing the Ct value to the standard curve (42). The stability and potential of the reference genes were evaluated using both GeNorm and NormFinder (GenEx, MultiD, Sweden) based on the Pfaffl equation (43).

### Statistics and relative quantification analysis

The threshold cycle (Ct) values of the gene of interest were normalised to that of the reference gene. The average values calculated were used for relative quantification of the gene of interest. The values obtained for transcript levels were used as a calibrator to determine whether a significant change in expression has occurred. Relative quantification levels were determined with the GenEx software (MultiD, Sweden). The equation describes one sample as the ratio of the gene of interest (target) versus a calibrator sample (control) and the reference gene (reference) versus a calibrator sample (control). The amplification efficiencies (E) were calculated as: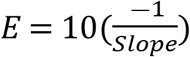. The difference in Ct values of the target gene in the control and sample (ΔCt target) and in the reference gene in the control and sample (ΔCt reference) were considered (43). The equation used to calculate the ratio was:

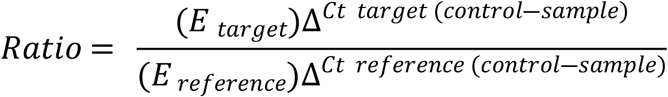

The experiments were carried out in four biological and three technical replicates. The relative expression level profiles of genes correlated with production of aflatoxins by the isolates were log transformed and assessed. Data analyses were made using GraphPad Prism version 5.02 (GraphPad Software, Inc., USA), One-way analysis of variance (1-way ANOVA), Tukey’s Multiple Comparison Test (TMCT), Post-test for linear trend and R statistical software (www.r-project.org), version 3.2.5.

## Acknowledgement

This work was supported by the University Science, Humanities and Engineering Partnerships in Africa (USHEPiA) Fund and South African Bio-Design Initiative (SABDI) grant number 420/01 SABDI 16/1021 secured by Dr NA Feto. Also, the authors acknowledge the University of Nairobi, Kenya and the University of Cape Town and Vaal University of Technology, South Africa for providing laboratory space and funding.

## Authors’ contribution

Authors, **AM, NAF, MSR**: conceived and designed the experiment, **AM:** generated and analysed the data, and drafted the manuscript. **NAF**: edited and reviewed the entire manuscript. **NAF, SO**, and **MSR**: reviewed the entire manuscript.

## List of figures

Figure S1. A flow diagram showing clustered genes (**a**) and the aflatoxin biosynthetic pathway (**b**). The corresponding enzymes and genes that are concerned with each step in the bioconversion are indicated in panel (**a)**. The vertical line gives a representation of the 82-kb aflatoxin biosynthesis pathway gene clusters and utilization of the sugar gene cluster in *Aspergillus flavus*. The novel gene names are represented on the left of the vertical line and the ancient gene names are shown on the right side The enzymes involved: fatty acid synthase, polyketide synthase, norsolorinic acid reductase, versiconal hemiacetal acetate reductase, esterase, versicolorin B synthase, versiconyl cyclase, desaturase, O-methyltransferase (MT-II), O-methyltransferase, O-methyltransferase (MT-I); AFB1: aflatoxin B1; AFB2: aflatoxin B2; AFG1: aflatoxin G1; AFG2: aflatoxin G2 (Adopted from(44–46). Asterisks (red star) represents the specific genes studied.

## Tables

Table S1 Integrity and quality of RNA assessed on Nano Drop spectrophotometer used for downstream analysis.

